# Specific ACE2 Expression in Cholangiocytes May Cause Liver Damage After 2019-nCoV Infection

**DOI:** 10.1101/2020.02.03.931766

**Authors:** Xiaoqiang Chai, Longfei Hu, Yan Zhang, Weiyu Han, Zhou Lu, Aiwu Ke, Jian Zhou, Guoming Shi, Nan Fang, Jia Fan, Jiabin Cai, Jue Fan, Fei Lan

## Abstract

A newly identified coronavirus, 2019-nCoV, has been posing significant threats to public health since December 2019. ACE2, the host cell receptor for severe acute respiratory syndrome coronavirus (SARS), has recently been demonstrated in mediating 2019-nCoV infection. Interestingly, besides the respiratory system, substantial proportion of SARS and 2019-nCoV patients showed signs of various degrees of liver damage, the mechanism and implication of which have not yet been determined. Here, we performed an unbiased evaluation of cell type specific expression of ACE2 in healthy liver tissues using single cell RNA-seq data of two independent cohorts, and identified specific expression in cholangiocytes. The results indicated that virus might directly bind to ACE2 positive cholangiocytes but not necessarily hepatocytes. This finding suggested the liver abnormalities of SARS and 2019-nCoV patients may not be due to hepatocyte damage, but cholangiocyte dysfunction and other causes such as drug induced and systemic inflammatory response induced liver injury. Our findings indicate that special care of liver dysfunction should be installed in treating 2019-nCoV patients during the hospitalization and shortly after cure.

## Introduction

In December 2019, a novel coronavirus, temporally named as 2019-nCoV by the World Health Organization (WHO), was discovered in Wuhan, Hubei Province, China [1]. The novel coronavirus was isolated from the airway epithelial cells of patients with pneumonia that was epidemiologically linked to Huanan seafood market in Wuhan. Within 2 months, 2019-nCoV spreads rapidly from Wuhan to all provinces in China, and more than 20 countries around the globe. As of 20:00 on February 3, 2020, 17,335 cases were confirmed in China, 21,559 were suspected, with an alarming number of 361 fatalities. Unfortunately, the numbers of both infected patients and fatalities are still growing rapidly with no effective drugs clinically approved. 2019-nCoV is closely related to Severe Acute Respiratory Syndrome coronavirus (SARS-CoV), and was proposed to share the same receptor, angiotensin converting enzyme 2 (ACE2), with SARS [2-4].

The liver, being one of the largest organs of the body in vertebrates, is located between the portal and the systemic circulation. It is constantly exposed to dietary antigens, viruses, and bacterial products with inflammatory potential. The liver damage can be caused by multitudinous factors such as exposure to toxins, excessive alcohol consumption, bile duct prevention obstruction, and viral infections [5]. Hepatitis B virus (HBV) is the main causative viral agent for hepatocellular carcinoma (HCC) in China [6]. A recent study also found acute infection of hepatitis E virus (HEV) causes varying degrees of liver damage [7]. Liver damage was reported in SARS-infected patients [8]. A study in 2015 has revealed that Middle East respiratory syndrome coronavirus (MERS-CoV) infection occurred in South Korea caused the rise of liver enzymes, implying liver function damage in patients infected with MERS-CoV [9]. An very recent epidemiologic study revealed that 43 out of 99 initial patients infected with 2019-nCoV had various degrees of liver function abnormality, with alanine aminotransferase (ALT) or aspartate aminotransferase (AST) above the normal range, and more important, one patient had severe liver function damage (ALT 7590 U/L, AST 1445 U/L) [1]. Being the most frequent organ damage out of the respiratory system, it’s currently unclear the liver damage of 2019-nCoV patients is caused directly by the viral infection of liver cells or by the drug toxicity. Previous RNA-seq data in the human protein atlas (HPA) database [10] shown relatively low expression of ACE2 in liver, as well as lung, which is the main target organ for 2019-nCoV and SARS. A low throughput study of ACE2 protein expressions in selected cell types of multiple organs has shown a low frequency of ACE2 occurrence in cholangiocytes, but not in hepatocytes, Kupffer cells, and endothelial cells [11], however the antibody detection might be subjected to non-specificity issue. Neither data sources could provide a definitive conclusion of cell type specific expression of ACE2 gene in liver. Recent advances of single cell technologies allow unbiased profiling of all cell types in given tissues at an unprecedented scale. To investigate the possible cause of 2019-nCoV patient liver damage, we used both published liver cell atlas data and an independent unpublished data generated in-house to evaluate ACE2 gene expression in all cell types in healthy livers.

## Methods

### Public dataset acquisition and processing

Gene expression matrix and cell type annotation of scRNA-seq data of normal human liver were downloaded from the Gene Expression Omnibus (GEO124395). We used Seurat 2.3 FindVariable function to select the top 2000 variable genes and performed principle component analysis. The first 20 principle components were used to project the data by tSNE. Cell labels provided by the authors were used for cell type annotations.

### Liver sample acquisition

Fresh normal liver tissues were taken from 4 deceased donors of liver transplants with an average age of 41. All cases donated organs after circulatory death (heart stop) complied with WHO regulations. Ethical approval was obtained from the research ethics committee of Zhongshan Hospital, Fudan University, and written informed consent was obtained from each patient.

### Tissue dissociation and single cell suspension preparation

Fresh tissue samples were collected and immediately stored in the GEXSCOPE Tissue Preservation Solution (Singleron Biotechnologies) at 2-8°C. Prior to tissue dissociation, the specimens were washed with Hanks Balanced Salt Solution (HBSS) for three times and minced into 1–2 mm pieces. The tissue pieces were digested in 2ml GEXSCOPE Tissue Dissociation Solution (Singleron Biotechnologies) at 37°C for 15min in a 15ml centrifuge tube with continuous agitation. Following digestion, a 40-micron sterile strainer (Corning) was used to separate cells from cell debris and other impurities. The cells were centrifuged at 1000 rpm, 4°C, for 5 minutes and cell pellets were resuspended into 1ml PBS (HyClone). To remove red blood cells, 2 mL GEXSCOPE Red Blood Cell Lysis Buffer (Singleron Biotechnologies) was added to the cell suspension and incubated at 25°C for 10 minutes. The mixture was then centrifuged at 1000 rpm for 5 min and the cell pellet resuspended in PBS. Cells were counted with TC20 automated cell counter (Bio-Rad).

### Single cell RNA sequencing library preparation

The concentration of single-cell suspension was adjusted to 1×10^5^ cells/mL in PBS. Single cell suspension was then loaded onto a microfluidic chip and single cell RNA-seq libraries were constructed according to the manufacturer’s instructions (Singleron GEXSCOPE Single Cell RNAseq Library Kit, Singleron Biotechnologies). The resulting single cell RNA-seq libraries were sequenced on an Illumina HiSeq X10 instrument with 150bp paired end reads.

### Primary analysis of scRNA-seq raw sequencing data

Raw reads were processed to generate gene expression matrices using an internal pipeline. Briefly, reads without poly T tails at the intended positions were filtered out, and then for each read cell barcode and unique molecular identifier (UMI) were extracted. Adapters and polyA tails were trimmed before aligning read two to GRCh38 with ensemble version 92 gene annotation. Reads with the same cell barcode, UMI and gene were grouped together to generate the number of UMIs per gene per cell. Cell number was then determined based on the inflection point of the number of UMI versus sorted cell barcode curve.

### Quality control, cell type clustering and visualization

To retain high quality cells, we removed cells with less than 200 genes or more than 5000 genes, as well as cells with mitochondrial content higher than 20%. 11550 cells were kept for the following analysis. The tSNE projection of the data was calculated similar to previous mentioned. Seurat FindClusters function was applied to obtain cell clusters with resolution 0.8. Cell types were assigned based on their canonical markers.

## Results

To assess the cell type specific expression of ACE2,, we first analyzed a published dataset containing scRNA-seq data of the normal liver samples from 9 colorectal patients (GEO124395) [12]. We divided single cells into subclusters based on the canonical markers and cell classification in the original literature (Figure 1A, B and C), and found specific ACE2 expressions in cholangiocytes and hepatocytes, though relatively sparse. In contrast, ACE2 expression was not observed in immune cells and stromal cells (Figure 1D). The observation partially agrees with previous data based on protein expression [11], that Kupffer cells and endothelial cells in liver are ACE2 negative. However, scRNA-seq identified low frequency of ACE2 expression in hepatocytes instead of negative for IHC.

**Figure 1.**
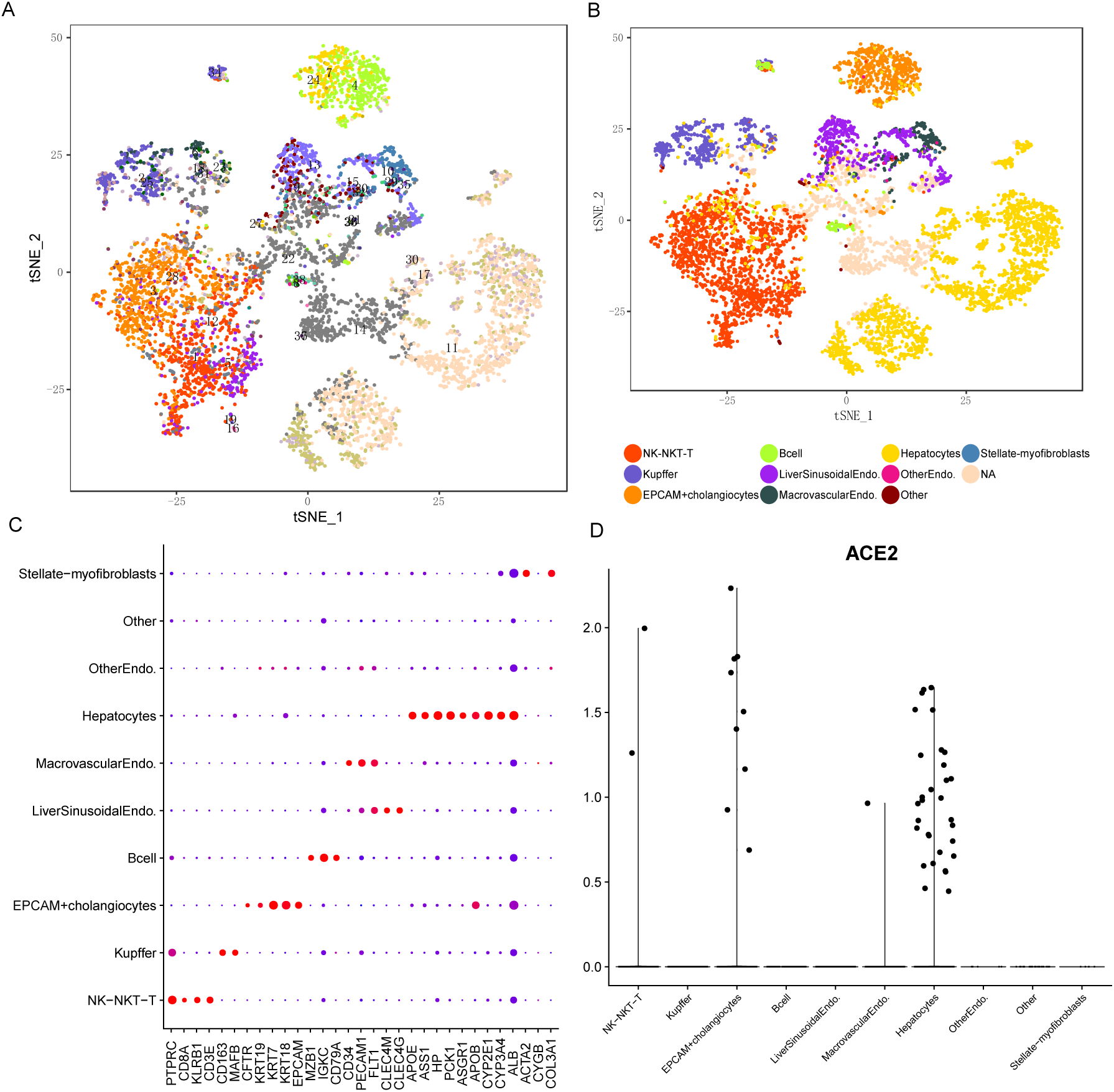
single cell analysis of published liver cell atlas. A) liver cell atlas visualized by tSNE, colored by cluster number. Cluster number information is provided by the authors. B) liver cell atlas visualized by tSNE, colored by major cell type compartments. Clusters were collapsed into major cell types based on their canonical markers. C) The dotplot showing cell type marker expression of all cell types. D) The violin plot of ACE2 gene expression of all major cell types.

To confirm the results, we further analyzed scRNA-seq data of an independent in-house cohort of 4 healthy donors (See Methods). Data were acquired from 11550 cells, and 16 cell types were identified, including parenchymal cell subpopulations and multiple non-parenchymal cell (NPC) subpopulations from the 4 patients (Figure 2A and B). For the parenchymal cells, hepatocytes and cholangiocytes were identified by well-known markers ALB and KRT19 as well as stem cell marker CD24 and SOX4.We also identified leucocytes (PTPRC) by their classical lineage markers (Figure 2C), for example, CD3D/E/G for T lymphocytes; killer Cell lectin like receptors KLRD1 and KLRF1 as well as NKG7 in CD3 negative population for Natural killer (NK) cells; MS4A1, CD79A, and HLA-II molecules for antigen presenting B cells; and MZB1, JCHAIN and IGHG1 for plasma cells. For mononuclear phagocytes (MP) populations, Kupffer cells (KCs) (MARCO, CD163, VSIG4), monocytes (CD14, FCN1, VCAN), and conventional dendritic cells (cDCs) (CD1C, FCER1A, HLA-II) were annotated by their respective markers. In addition, neutrophils were identified by expression of specific marker of CSF3R, G0S2, CXCR2 and FCGR3B. We also confirmed a granulocyte progenitor cell cluster (MPO, ELANE, AZU1) and a small cluster of erythrocytes (HBA1, HBB). Other NPC types like endothelial cells and myofibroblasts were distinguished by pan-endothelial marker CDH5 and myoid marker ACTA2 and MYH11. Endothelial cells were further classified as liver sinusoidal endothelial cells (LSEC) and macrovascular endothelial cells (MaVEC) based on the different signatures [12].

**Figure 2.**
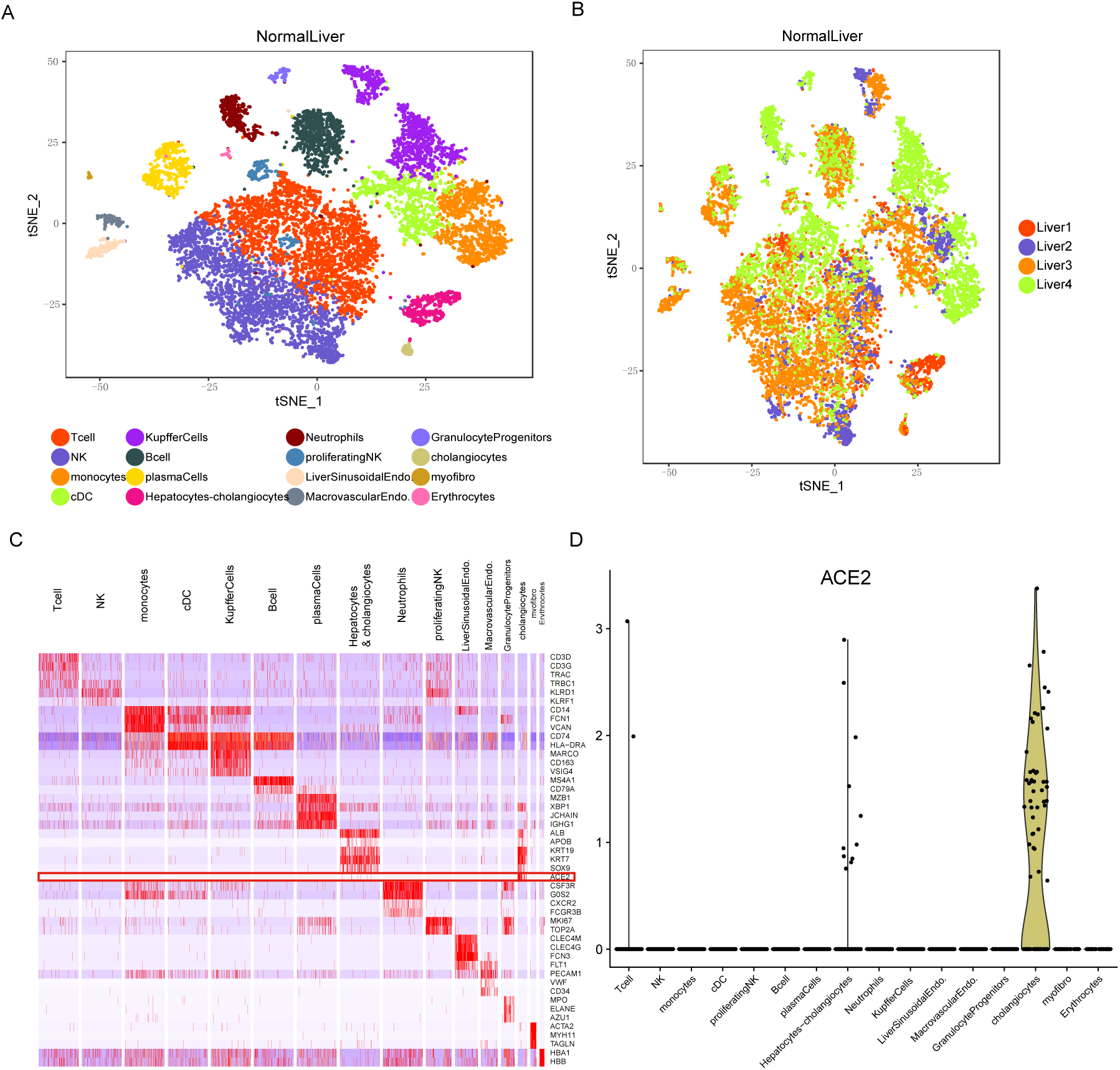
single cell analysis of the in-house cohort of healthy livers. A) Cell type annotations of all four patients by the tSNE plot. Hepatocyte-cholangiocytes cluster includes potential stem-like cells capable of differentiating into both cell types. B) tSNE plot colored by patients C) Heatmap of marker genes of 16 cell types identified. The red box highlights the cell type specific expression of ACE2. D) The violin plot of ACE2 gene expression across all cell types.

We found significant enrichment of ACE2 expression in a major portion of the cholangiocytes cluster (59.7% of cells). Similar to the public dataset analyzed above (Figure 1D), the results from this cohort confirmed that immune and stromal compartments were ACE2 negative (Figure 2D). We observed low expression of ACE2 in hepatocytes (2.6%), with average expression level 20 fold less than the expression level in the cholangiocytes population. Significantly, but different from the protein analysis, ACE expression level in cholangiocytes is comparable to Alveolar Type 2 cells [13, 14], which is the major SARS and 2019-nCoV targeting cell type in lung. Therefore, ACE2 expression in cholangiocytes may suggest a potential mechanism of infection and direct damage of bile ducts by virus using ACE2 as host cell receptors, such as 2019-nCoV and SARS.

## Discussion and Conclusion

Based on the scRNA-seq data of two independent cohorts, we have identified cholangiocyte-specific expression of ACE2 in healthy human livers and normal liver samples from colorectal cancer patients. Since ACE2 is capable in mediating 2019-nCoV and SARS infections, its expression pattern reveals possibility for direct infection of cholangiocytes by 2019-nCoV and SARS. However, our findings suggest that hepatocytes might not be targeted by these viruses, or at least not through ACE2. Therefore, we speculate that liver damages in patients infected by 2019n-CoV might not be caused directly by viral infection of hepatocytes. Cholangiocytes are multifunctional and play critical roles in liver regeneration and immune responses [15]. Thus, the potential damage of cholangiocytes by 2019-nCoV may lead to profound consequences in the liver. Our findings also suggest that the liver abnormality of 2019-nCoV patients may be caused by drug used in the treatment, or systemic inflammatory response-induced by pneumonias. We could not exclude the possibility of technical limitations in detecting extremely low ACE2 expression in hepatocytes at this point.

This study demonstrated the highly sensitive nature of single-cell resolution analysis and facilitated the understanding of the mechanisms of liver malfunction in 2019-nCoV-infected patients. Such information call for patient cares regarding liver responses, especially related to cholangiocyte function, of the large number of 2019-nCoV patients currently under emergency and potential post-cure treatment for liver recovery after hospitalization.

## Tables

**Table 1.** Patient information of the in-house cohort.

| Sample ID | Age | Gender |
| --- | --- | --- |
| Liver1 | 36 | M |
| Liver2 | 41 | M |
| Liver3 | 44 | M |
| Liver4 | 42 | F |

